# Inner-ear delivery of AAV2-retro robustly labels lateral olivocochlear efferents but results in off-target transfection in the contralateral cochlea and the brain

**DOI:** 10.64898/2025.12.03.692215

**Authors:** Mahmoud Khalil, Mikayla Bisignani, Karl Kandler

## Abstract

Gene delivery via adeno-associated viruses (AAVs) is a valuable tool for understanding the organization and function of auditory neuronal circuitry. In this study, we explored the use of AAVs injected into the inner ear to transfect the lateral olivocochlear (LOC) pathway, a poorly understood ipsilateral component of the auditory efferent feedback system. AAV serotypes exhibit a wide range of properties, including cell type selectivity and the capacity for retrograde transport, but none of the AAV serotypes used for inner ear delivery have been reported to successfully transfect the LOC pathway. Here, we show that unilateral inner ear delivery of AAV2-retro robustly labeled the LOC pathway. However, the ability of AAV2-retro to diffuse via the cochlear aqueduct from the perilymph into the cerebrospinal fluid resulted in off-target transfection in the brain and the labeling of LOC projections in the contralateral cochlea. The extent of contralateral transfection of LOC neurons depended on the age of the animal and the amount of virus delivered to the inner ear. These findings highlight the importance of considering the dosage and properties of AAV serotype when interpreting experimental results.

## Introduction

Adeno-associated viruses (AAVs) are an indispensable tool for neuroscience research with applications ranging from circuit tracing (Luchicchi et al, 2021; Saleeba et al, 2019; Weinholtz et al, 2020) and circuit manipulation (Watakabe et al, 2017) to gene therapy (Deverman et al, 2018). In the field of auditory neuroscience, inner ear delivery of AAVs has been used to develop potential therapeutic approaches to rescuing hearing loss (Hao et al 2019). AAV-mediated gene therapy has been successfully applied to restore hearing in congenital deaf mice delivering functional genes to hair cells (Pan et al, 2017; Akil et al, 2012) or through the delivery of miRNA to rescue autosomal dominant hearing loss (Shibata et al, 2016).

Different serotypes of AAV vary in their tropism for different cell types (Aschauer et al, 2013; Sun et al, 2019; Witteveen et al, 2024), their capacity for anterograde and retrograde transport (Rothermel et al, 2013; Wang et al, 2021(A)), their efficacy of viral transduction (Aschauer et al, 2013), and their local diffusion from the injection site (Taymans et al, 2007; Witteveen et al, 2024). Recent studies have identified AAV variants that are suited for AAV-mediated gene delivery in the inner ear by developing variants that exhibit lower toxicity and higher efficacy of transfecting hair cells and supporting cells in the injected ear (Ivanchenko et al, 2021; Landegger et al, 2017; Isgrig et al, 2019)

In this study, we tested the suitability of inner ear delivery of AAVs to retrogradely transduce the efferent lateral olivocochlear (LOC) pathway. The LOC pathway and the medial olivocochlear (MOC) pathway are the two components of efferent innervation in the mammalian cochlea (Warr et al, 1979; Guinan et al, 1983; White et al, 1983). LOC neurons predominantly project ipsilateral across species, including mice (Campbell and Henson, 1988; Maison et al, 2002; Brown et al 1993, 2008), rats (White and Warr, 1983; Warr et al, 1997; Abou-Madi et al, 1987; Cantos et al, 2000; Aschoff et al, 1988; Horvath et al, 2000; Vetter et al, 1992), Guinea pigs (Niu et al, 2004; Horvath et al, 2003), Bats (Aschoff et al, 1987; Bishop et al, 1987), hamsters (Sánchez-González et al, 2003), gerbils (Kaiser et al, 2011; Aschoff et al, 1988), chinchillas (Azeredo et al, 1999), monkeys (Thompson et al, 1986), and cats (Warr et al, 1979; Guinan et al, 1983). LOC neurons can be subdivided into dopaminergic shell LOC neurons which are located around the boundary of the lateral superior olive (LSO) and cholinergic core LOC neurons which are located inside the LSO nucleus (Darrow et al, 2006; Warr et al, 1997; Brown, 1987). Core LOC neurons represent 80-95% of all LOC neurons depending on species (Darrow et al, 2006; Horváth et al, 2000; Vetter et al, 1992). Both core and shell LOC neurons innervate the afferent synapses at the inner hair cell region (Wilson et al, 1991; Liberman, 1980), with core LOC projections forming dense arborizations along tonotopically restricted regions and shell LOC projections bifurcating along the cochlear axis to cover a wide tonotopic region with en passant swellings instead of dense terminal arbors (Warr et al 1997; Brown et al, 1987). Additionally, the efferent projections of core LOC neurons are almost exclusively ipsilateral, while 5-20% of shell LOC neurons also project to the contralateral cochlea (Horváth et al, 2000; Vetter et al, 1992).

Previous studies suggest that the LOC pathway plays a role in protecting against both age- and noise-induced synaptopathy (Liberman et al, 2014; Darrow et al, 2007), in learning (Irving et al, 2011), and in balancing interaural inputs (Darrow et al, 2006). In humans, the function of LOC efferents was speculated to be involved in the balancing of binaural input in tinnitus patients (Tae et al 2022) and in auditory symptoms associated with Parkinson’s disease (Pisani et al, 2015). However, a better understanding of LOC pathway function and its involvement in pathologyhas been hampered by the lack of a biological readout specific to LOC pathway activity and the difficulty of selectively targeting LOC neurons without impacting other components of the auditory system. Anatomical lesions of either the LSO nucleus, where LOC neurons reside (Darrow et al 2006; Liberman et al 2014), or the olivocochlear bundle that contains LOC axons (Irving et al 2011) are not specific to LOC neurons. Additionally, the shared cholinergic identity between most MOC and LOC neurons limits avenues for pharmacological manipulations. Targeting the LOC pathway through inner ear delivery of AAV could open up additional approaches to overcome these hurdles. However, AAV serotypes commonly used for gene delivery to the inner ear have not been reported to label the LOC pathway (Kilpatrick et al, 2011; Blanc et al, 2022; Marcovich et al, 2022). Identifying a suitable AAV serotype for transducing the LOC pathway would, therefore, be a promising first step. A major challenge in identifying a suitable serotype, however, is that the capacity for retrograde transport and selectivity for neuronal subtypes varies widely among AAV serotypes (Rothermel et al, 2013; Wang et al, 2021(A)).

AAV2-retro is an AAV variant capable of retrograde neuronal transduction that was created by modifying the capsid of the naturally occurring AAV2 serotype (Tervo et al, 2016). AAV2-retro exhibits variable cellular tropism, however, as it shows diminished retrograde transduction capacity for some neuron types (Han et al, 2023; Skorput et al, 2022; Wang et al, 2018) while failing to transduce others (Bortoloci et al, 2022). When delivered to the inner ear, AAV2-retro was observed to transduce MOC and shell LOC neurons but not core LOC neurons (Wang et al, 2021 B). While this observation could be reflective of AAV2-retro’s tropism, successful AAV transduction could also be impacted by other factors like the specific promoter used to drive gene expression (Kügler et al, 2003; Bohlen et al, 2020) and the titer used (Lu et al, 2024). Since AAV2retro is one of the most readily available AAV serotypes capable of robust retrograde transduction, we set out to investigate whether it can retrogradely transduce core LOC neurons. Our results demonstrate that inner ear delivery of AAV2-retro results in robust retrograde transduction of core LOC neurons in both neonatal and juvenile mice. Additionally, we report that unilateral delivery of AAV2-retro results in bilateral transduction of core LOC neurons through the diffusion of viral particles through the cochlear aqueduct into the cerebrospinal fluid and the contralateral ear. The extent of AAV2-retro diffusion throughout the central nervous system was volume- and age-dependent. Taken together, our results demsonstarte that AAV2-retro is a viable method of retrogradely transducing the LOC pathway but that its proclivity to diffuse throughout the central nervous system should be taken into account when designing experiments and interpreting results.

## Materials and methods

### Animals

All experiments were performed on C57Bl6/J mice of both sexes. All experimental procedures were in accordance with the NIH guidelines and were approved by the institutional animal care and use committee at the University of Pittsburgh.

### Inner ear delivery of AAV and Fast Blue

Surgeries for inner ear delivery of AAV were performed on animals aged postnatal day (P) 1-3 and P21-28. In neonates, r-AAV2-retro-hSyn-mCherry (AAV2-reto; Addgene 114472-AAVrg; titer 2e+13 GC/mL) or Fast Blue (Polysciences inc, catalogue number: 17740) was delivered to the inner ear through injections into the posterior semicircular canal (Isgrig et al 2018). P2-3 mouse pups were anesthetized using hypothermia until the cessation of movement and a lack of response to tail pinch. Hypothermic anesthesia was maintained for the duration of the surgery by placing the pup on ice wrapped with a sterile absorbent pad. An incision was made behind the ear flap to expose the sternocleidomastoid muscles, which were dissected to expose the cartilaginous structure of the posterior semicircular canal. AAV was injected into the semicircular canal with a pulled glass micropipette attached to a stereotaxic mount. Injections were made at a rate of 1 nl/s (Nanoject III) or with pulses of 2 nl at 23nl/s every three seconds (Nanojet II). Total volume injected in neonates was 360-400 nl for Figure 1, and 200 nl, or 50 nl as indicated in later figures. After completing the injection, the pipette was removed, and the incision was sutured with an 8/0 monofilament nylon suture (Fine Science Tools). Pups were then placed on a warming pad and returned to its mother once it recovered from anesthesia and gained mobility.

**Figure 1:**
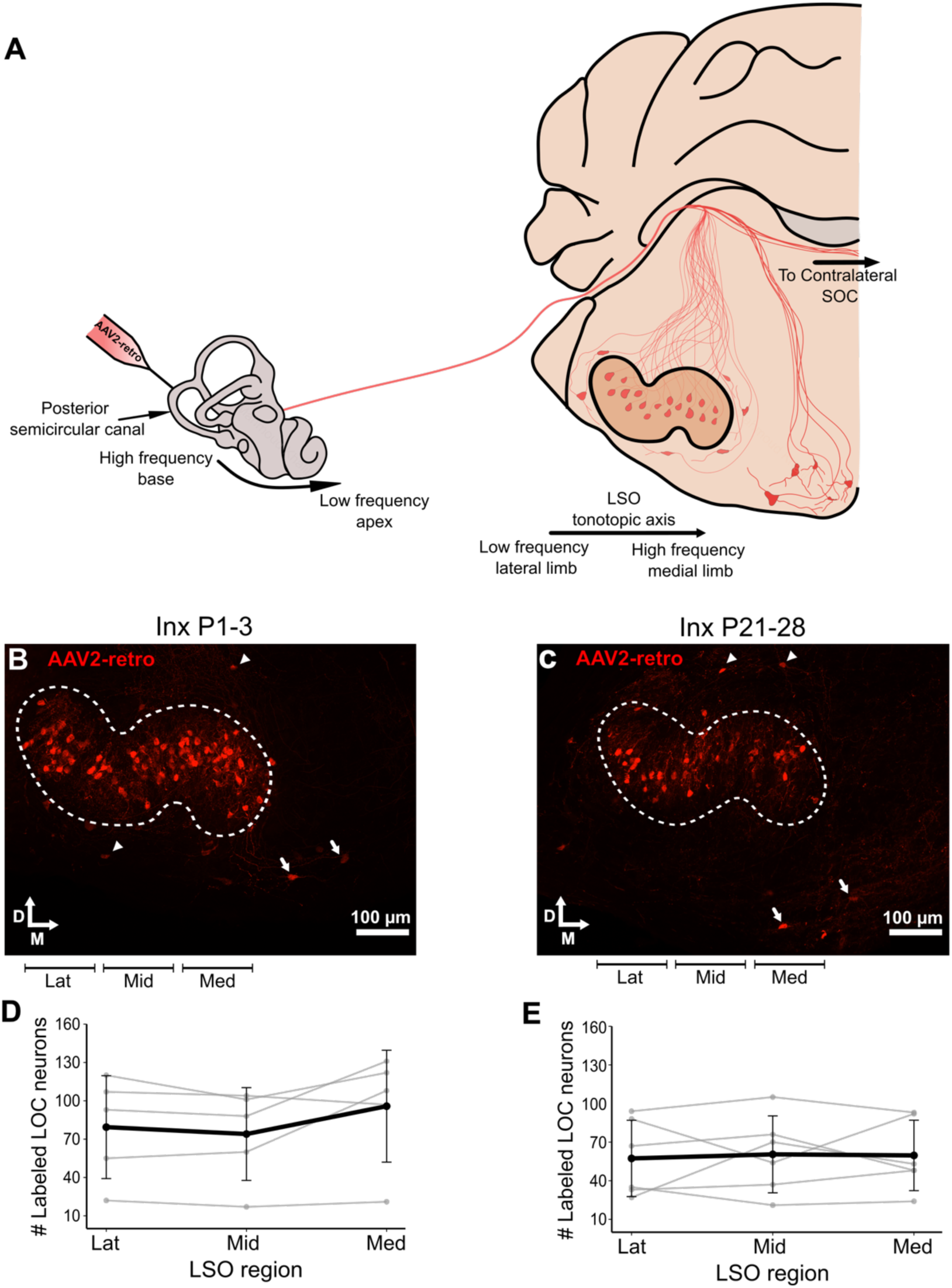
Inner ear delivery of AAV2-retro robustly labels LOC neurons along the tonotopic axis of the LSO. A) Overview of the lateral olivocochlear pathway and experimental approach. B) Retrogradely labeled core LOC neurons in the ipsilateral LSO following neonatal inner ear delivery of AAV2-retro. D= dorsal. M= medial. Arrowheads: Shell LOC neurons. Arrows: MOC neurons, Lat: Lateral. Mid: Middle. Med: Medial. Inx: Injection C) Retrogradely labeled core LOC neurons in the ipsilateral LSO following inner ear delivery of AAV2-retro in P21-28 animals. D) Quantification of retrogradely labeled LOC neurons in the lateral, middle, and medial regions of the ipsilateral LSO following P1-3 inner ear delivery of AAV2-retro (N = 5). Thin line – individual cases, thick line average. Lateral region: 79±40, middle region: 74±36, medial region: 95±43. oneway repeated measures ANOVA, F=2.76 P=0.12. E) Quantification of retrogradely labeled LOC neurons in the lateral, middle, and medial regions of the ipsilateral LSO following P21-28 inner ear delivery of AAV2-retro (N = 6). Lateral region: 57±29, middle region: 59±27, medial region: 60±29 one-way repeated measures ANOVA, F=0.064 P=0.937.

P21 and older mice were anesthetized by intraperitoneal injection of 100 mg/kg ketamine and 10 mg/kg xylazine. AAV injections into the posterior semicircular canal were similar to Suzuki et al, (2017). A postauricular incision was made and a small incision through the underlying lipid and connective tissue exposed the sternocleidomastoid muscles, which were dissected in order to expose the posterior semicircular canal. A small hole was hand-drilled in the semicircular canal using a 27-gauge hypodermic needle. Perilymph was allowed to drain for 5-10 minutes and was soaked up using a sterile Gelfoam sponge (Pfizer injectables, manufactured and distributed by Pharmacia & Upjohn). A 0.0030’’/0.0040’’ ID/OD Polyimide catheter tube (manufactured by Teleflex Medical OEM) attached to a glass pipette with UV curable epoxy was then inserted into the posterior semicircular canal. The opening through which the polyimide catheter was inserted was sealed using fragments of muscle tissue and cyanoacrylate glue (3M Vetbond Tissue Adhesive). Injections were performed at a rate of 1 nl/s (Nanojet III) or with pulses of 23 nl at 23nl/s every 20 seconds (Nanojet II). Total injection volume in juvenile mice was 1.2 microliters. After withdrawal of the catheter, the hole in the semicircular canal was sealed with muscle fragments and cyanoacrylate glue (3M Vetbond Tissue Adhesive). The surgery site was sutured with a 5-0, FS-2, 19 mm reverse cutting, Perma-Hand Silk suture (Ethicon). Lidocaine cream was applied over the sutured incision to remedy pain, and for three days animals received daily subcutaneous injections of 10mg/kg ketoprofen.

### Immunocytochemistry for AAV and Fast Blue tracing experiments

21-22 days after AAV injections and 14 days after Fast Blue injections, animals were anesthetized with 100mg/kg ketamine and 10mg/kg xylazine and perfused transcardially with phosphate buffered saline (PBS) followed by 4% paraformaldehyde in PBS (4% PFA). The brains and temporal bones were then dissected and post-fixed in 4% PFA in PBS solution overnight at 4 °C followed by submersion in 30% sucrose in PBS for cryopreservation for 48 hours at 4 °C. Coronal sections (50 µm thick) of the superior olivary complex (SOC) were prepared using a freezing microtome (Microm HM 430). Temporal bones underwent cochlear whole mount dissections using fine forceps (Haque et al, 2015).

Brain sections from animals injected with Fast Blue were washed in PBS (2x for 5 min) in PBS and were mounted on superfrost plus microscope slides (Fisherbrand) using Fluoromount-G (Southern Biotech) and coverslipped.

Brain sections and cochlea tissue from AAV tracing animals were washed in PBS (2x for 5 min) and then incubated in blocking solution (1% BSA, 5% Donkey serum, 0.1% triton-x) overnight at 4 °C. Both primary and secondary antibodies were prepared in blocking solution. The following antibodies were used 1:500 Scigen Goat anti-Tdtomato (Lot#: 0081030221) and 1:1000 Invitrogen Alexa Fluor 568 donkey anti goat (Ref#: A11057, Lot#:2304269).

Following blocking, brain sections and cochlea tissue were incubated in primary antibody solution overnight at 4 °C on a stirrer plate, washed in PBS (5x for 10 min), and incubated in secondary antibody solution overnight at 4 °C on a stirrer plate. Sections were washed in PBS (5x for 10 min), mounted on superfrost plus microscope slides (Fisherbrand) using Fluoromount-G (Southern Biotech) and coverslipped.

### DiI tracing

P2-3 mouse pups were anesthetized with 100mg/kg ketamine and 10mg/kg xylazine and transcardially perfused with PBS followed by 4% paraformaldehyde in PBS (4% PFA). An incision was made ventro-caudal to the ear flap, and the underlying tissue was dissected in order to expose the auditory bulla. Individual cotton threads separated from Curity gauze sponges (Coviden) infused with DiI were then inserted through the auditory bulla and into the cochlea using fine syringes. Threads were infused with DiI by soaking them with 1,1’-Dioctadecyl-3,3,3’,3’tetramethyl-indocarbocyannine perchlorate (DiI, Aldrich, Lot# MKCDO794) dissolved in 70% ethanol and subsequent air drying. Tissues were then stored in a 4% PFA/PBS solution in the dark at room temperature and after 14-15 weeks, the brains were extracted and sliced coronally (50 µm) in PBS using a Leica VT1000S Vibratome Sections containing the SOC were mounted on superfrost plus microscope slides (Fisherbrand) sing Fluoromount-G (Southern Biotech) and coverslipped.

### Microscopy and quantification of cells

Digital micrographs of the SOC and cochlea were obtained using a fluorescent microscope (Axio Imager, Zeiss) or a confocal microscope (Nikon A1). Labeled cells were counted manually using the “multi-point” tool on ImageJ version 2.9.0/1.54f. The identity of retrogradely labeled olivocochlear neurons was determined based on anatomical location. Core LOC neurons were defined as retrogradely labeled neurons located within the LSO. The lateral, middle and medial regions of the LSO were determined through dividing the LSO along the medi-lateral axis into three equal regions. Shell LOC neurons were defined as bi- and multi-polar cells located around the periphery of the LSO nucleus. MOC neurons were defined as retrogradely labeled cells located in the ventral nucleus of trapezoid body medio-ventral to the LSO.

### Statistical Analysis

Rstudio was used to conduct all the statistical analysis and generate plots. Plots were created using the tidyverse and ggplot2 packages. Data was tested for normalicy by the Shapiro–Wilk normality tests using the base R stats package. Statistical comparisons were performed using either Student’s t-tests or ANOVA (base R stats package). Tukey’s HSD post-hoc comparisons were performed if the ANOVA results were significant by using the emmeans package. The Holm–Bonferroni correction for multiple comparisons was performed to correct the P-values reported for figures 2DF and 2 J-L using the base R stats package.

**Figure 2:**
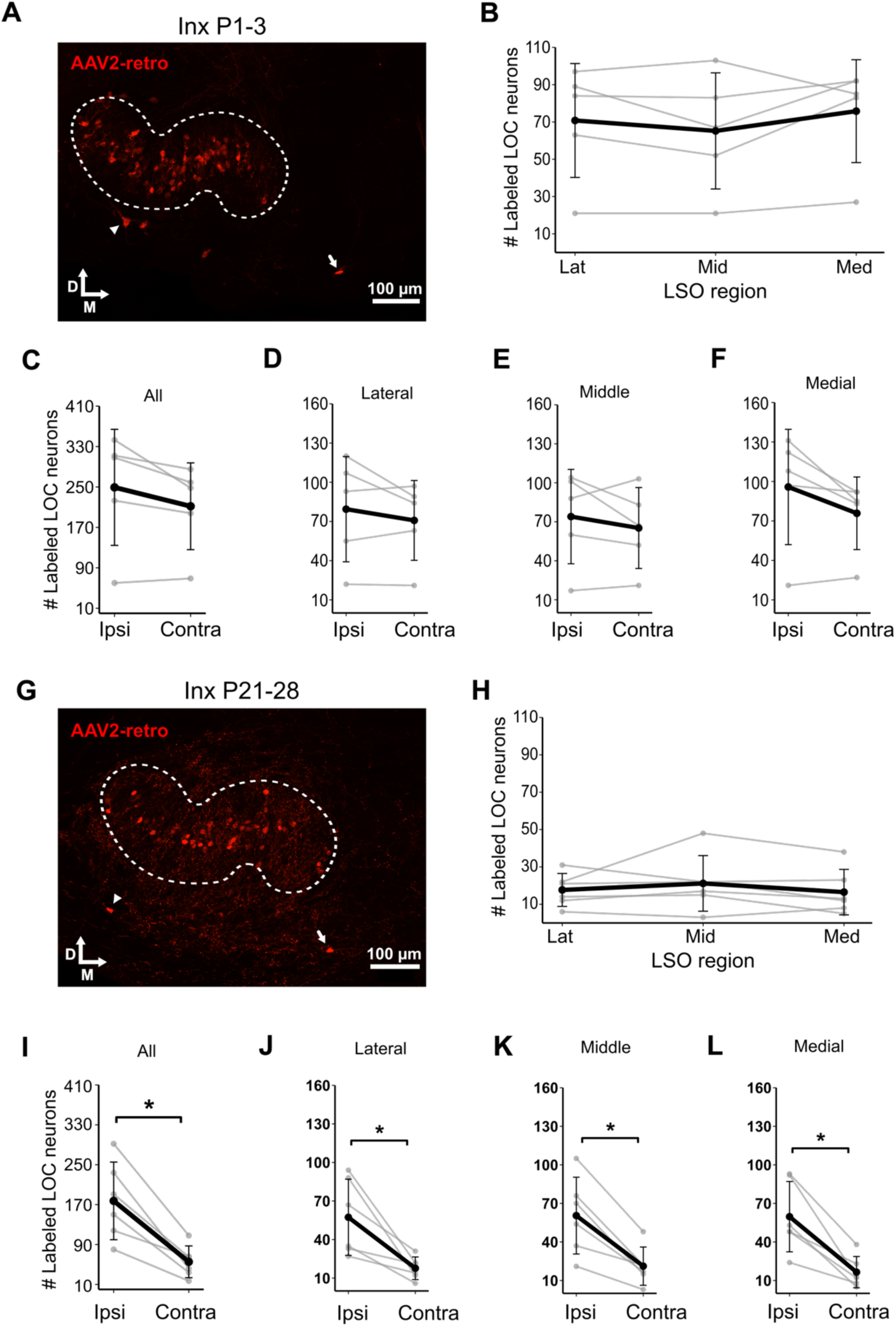
Unilateral inner ear injection of AAV2-retro labels LOC neurons in the contralateral LSO. A) Retrogradely labeled core LOC neurons in the contralateral LSO in neonatally injected mice. D= dorsal. M= medial. Arrowheads: Shell LOC neurons. Arrows: MOC neurons, Lat: Lateral. Mid: Middle. Med: Medial. Inx: Injection B) Quantification of retrogradely labeled LOC neurons in the lateral, middle, and medial regions of the contralateral LSO in neonatally injected mice. N = 5 lateral region: 71±30, middle region: 65±31, medial region: 76±28, one-way repeated measures ANOVA, F=2.76, P=0.12. C) Comparison of the overall number of retrogradely labeled LOC neurons between the ipsilateral and the contralateral LSO in neonatally injected mice. N = 5 Paired t-test, t=2.18, P=0.09. D-F) Region-specific comparisons between ipsilateral and contralateral LSO in neonatally injected mice. N = 5 Paired t-tests with HolmBonferroni correction for multiple comparisons. Lateral Region t=1.1 P= 0.66. Middle region t=1.01 P= 0.66. Medial region t=2.16 P= 0.288. G) Retrogradely labeled core LOC neurons in the contralateral LSO in juvenile mice injected with AAV2-retro. H) Quantification of retrogradely labeled LOC neurons in the lateral, middle, and medial regions of the contralateral LSO in juvenile mice injected with AAV. N = 6 lateral region: 18±9, middle region: 16±12, medial region: 21±15, one-way repeated measures ANOVA, F=0.65, P=0.54. I) Comparison of the overall number of retrogradely labeled LOC neurons between the ipsilateral and the contralateral LSO in juvenile mice injected with AAV2-retro. N = 6 Paired t-test, t=5.1, P=0.0038. J-L) Region-specific comparisons between ipsilateral and contralateral LSO in juvenile mice injected with AAV2-retro. N = 6 Paired t-test with Holm-Bonferroni correction for multiple comparisons. Lateral region t=3.44 P= 0.018. Middle region t=5.04 P= 0.01. Medial region t=4.67 P= 0.01.

## Results

### Inner ear injections of AAV2-retro robustly transfect LOC neurons in neonatal and juvenile mice

Following neonatal semicircular canal injection of AAV2-retro (Figure 1A), we observed robust retrograde labeling of MOC neurons, and extensive labeling of retrogradely transfected neurons in and around the ipsilateral LSO (Figure 1B). The large size and bi- or multipolar morphology of the neurons on the periphery of the LSO were consistent with a shell LOC identity (Warr et al, 1997; Darrow et al 2006; Wu et al 2020); whereas, the small, oval, and mostly fusiform morphology of the neurons positioned in the center of the LSO corresponded with the characterization of core LOC neurons in the literature (Campbell and Henson, 1988; Warr et al, 1997; Cantos et al, 2000; Brown et al, 2008; Vetter et al, 1992). The number of retrogradely labeled core LOC neurons was 249 ± 115 (N=5), which is in line with what has been reported previously for mice using retrograde tracing (112-307 neurons, Brown et al, 2008; Campbell and Henson, 1988) and with the number of core LSO neurons determined by immunohistochemistry using acetylcholinesterase as a marker (214 neurons, Brown et al, 2008). Our results thus indicate that a single injection of AAV2-retro into the PSCC of neonatal mice can label the entire population of ipsilateral core LOC neurons.

The projections of core LOC neurons are tonotopically organized (Guinan et al, 1984), with neurons located in the lateral regions of the LSO projecting to the low frequency apical regions of the cochlea and neurons located in the medial regions of the LSO projecting to the high frequency, basal regions of the cochlea. Some AAV serotypes exhibit reduced transfection efficacy of cochlear regions distal from their injection site (György et al, 2017; Yoshimura et al, 2018; Zhang et al 2023), while other serotypes exhibit varying levels of tropism to cells in the different regions of the cochlea (Iranfar et al 2025; Ivanchenko et al 2021). We quantified the number of retrogradely transfected LOC neurons in the corresponding lateral, middle, and medial regions of the LSO to determine whether or not AAV2-retro equally transfected LOC neurons along the entire length of the cochlea (Figure 1C). The medial region had the highest number of retrogradely labeled LOC neurons at 95±43 followed by the lateral region at 79±40 neurons and the middle region at 74±36 (N=5). The differences between the three regions, however, was not significant (one-way repeated measures ANOVA, F=2.76, P=0.12), which indicates that AAV2-retro is both capable of spreading from the posterior semicircular cana to all cochlear region and displays similar tropism LOC neurons across cochlear regions.

The neuronal tropism of several AAV serotypes decreases with age of the host animal (Chakrabarty et al 2013). In the mouse cochlea, the tropism of AAV-Anc80L65 and AAV-S for spiral ganglion neurons is higher in neonatal cochlea compared to later ages (Ivanchenko et al 2021; Richardson et al 2021). To investigate whether AAV2-retro still robustly transfects LOC neurons at older ages, we unilaterally delivered the viral vector to the inner ears of P21-28 mice. LOC neurons in the ipsilateral LSO of these mice were robustly transfected by AAV2-retro (Figure 1D), with an average of 178 ± 77 neurons (N=6), which is within the range previously reported in the literature (112-307 neurons, Brown et al, 2008; Campbell and Henson, 1988) and is similar to the number we observed in neonatally injected mice (249 ± 115 N=5; two-tailed t-test t= 1.407, P= 0.193). The number of retrogradely labeled LOC neurons in the lateral, middle, and medial regions of the ipsilateral LSO were similar at 57±29, 59±27, and 60±29 respectively (Figure 1E, N=6, one-way repeated measures ANOVA, F=0.064 P=0.937). These results indicate that AAV2-retro is an effective vector for retrograde transfection and gene delivery to core LOC neurons in both neonatal and juvenile mice.

### Unilateral inner ear delivery of AAV2-retro results in transfection of LOC neurons in the contralateral LSO

Previous studies using a wide variety of anatomical tracers have demonstrated that the projections of core LOC neurons are almost entirely ipsilateral in mice (Campbell and Henson, 1988 horseradish peroxidase; Brown et al, 1993 biocytin and horseradish peroxidase; Brown et al, 2008 fluorogold and horseradish peroxidase). In contrast to these studies, our unilateral inner ear injections of AAV2-retro labeled core LOC neurons in the contralateral LSO in both neonatal (Figure 2A) and juvenile (Figure 2G) mice. Neonatal mice had an average of 212 ± 85 retrogradely labeled LOC neurons in the contralateral LSO (N=5), which was similar to the 249 ± 115 neurons in the ipsilateral LSO (Figure 2C, paired t-test, t=2.18, P=0.09). Retrogradely transfected LOC neurons in the contralateral LSO were equally distributed along its mediolateral axis, with no significant differences observed between the lateral, middle, and medial regions (Figure 2B, oneway repeated measures ANOVA, F=2.76, P=0.12). Comparing the number of labeled LOC neurons in each of these LSO regions between the ipsilateral and contralateral LSO revealed no significant differences either (Figure 2D-F; paired t-test corrected for multiple comparisons; lateral P= 0.66; Middle P= 0.66; Medial P= 0.288).

In contrast to neonatal mice, in juvenile mice, the number of retrogradely labeled LOC neurons in the contralateral LSO 55±31, N=6) was significantly less than in the ipsilateral side (178 ± 77 neurons N=6; Figure 2G-I, Paired t-test, t=5.1, P=0.0038). The reduction in the number of labeled neurons in the contralateral LSO was not confined to a single isofrequency region as the distribution of the labeled neurons was uniform across the mediolateral axis (Figure 2H; one-way ANOVA F=0.65 P=0.54) and all three individual LSO regions had significantly less LOC neurons labeled when compared to their ipsilateral counterparts (Figure 2J-L; paired t-test corrected for multiple comparisons; lateral P= 0.018; Middle P= 0.01; Medial P= 0.01).

### AAV Injected into the Inner Ear Travels Through the Cerebrospinal Fluid and Into the Contralateral Inner Ear

The extensive contralateral labeling of core LOC neurons following unilateral inner ear delivery of AAV2-retro was surprising in light of ipsilateral nature of the LOC pathway reported earlier (Campbell and Henson, 1988; Brown et al, 1993; Brown et al, 2008). While contralateral labeling in juveniles was lower than in neonates, it remained substantial (~23%) and far above the 0–1% typically reported after unilateral tracer injections (Campbell and Henson, 1988; Brown et al, 1993; Brown et al, 2008). The perilymph of the cochlea is connected to the cerebrospinal fluid (CSF) of the central nervous system via the cochlear aqueduct (Kellerhals, 1979). This connection allows for the bidirectional flow of material (Talaei et al, 2019), including viral particles (Mathiesen et al 2023), between the perilymph and the CNS. We hypothesized, therefore, that the robust labeling of LOC neurons in the contralateral LSO was most likely mediated by the diffusion of viral particles from the injected inner ear to the CSF via the cochlear aqueduct and onto the contralateral cochlea. To test our hypothesis, we surveyed the brain of mice that underwent inner ear AAV2retro delivery for evidence of ectopic transfection of neurons that do not project to the cochlea and would have only been transfected by viral particles that diffused out of the cochlear perilymph into the CSF. Indeed, we observed ectopic transfection of cerebellar neurons in all the mice that underwent the inner ear delivery of AAV2-retro as neonates (Figure 3A) or as juveniles (Figure 3B).

**Figure 3:**
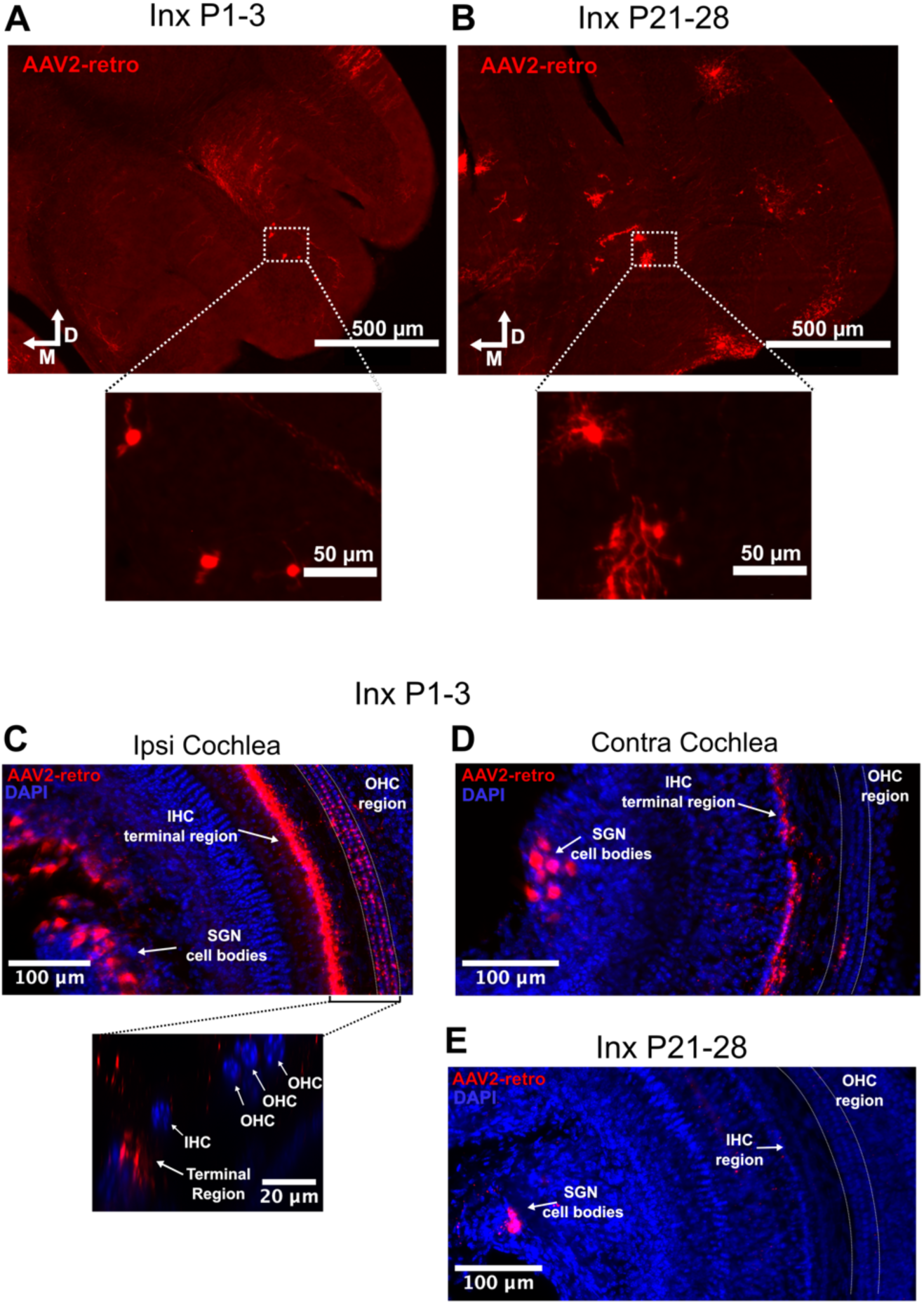
AAV2-retro injected into the inner ear diffuses throughout the cerebrospinal fluid and into the contralateral cochlea. A) Transfected cerebellar neurons following neonatal inner ear delivery of AAV2-retro. M= medial, D= dorsal. B) Transfected cerebellar neurons following juvenile inner ear delivery of AAV2-retro. C) cochlea from the injected ear following neonatal inner ear delivery of AAV2-retro along with a magnified view of the organ of Corti cross-section showing AAV labeling of terminal synapses underneath the inner hair cell region. D) Contralateral cochlea following neonatal inner ear delivery of AAV2-retro. E) Contralateral cochlea following juvenile inner ear delivery of AAV2-retro.

To further test our hypothesis, we examined the injected cochlear for direct somatic transfection by AAV2-retro and whether we could observe similar transfection in the contralateral cochlea. In the injected cochlea, AAV2-retro robustly transfected the cell bodies of spiral ganglion neurons (SGN) (Figure 3C) and we observed transfected SGN cell bodies also in the contralateral cochlea in both neonatal animals (Figure 3D) and juveniles (Figure 3E). Taken together, these results suggest a diffusion of viral particles from the injected inner ear to the contralateral cochlea, where it infects the terminals of efferent fibers to retrogradly label of LOC neurons in the contralateral LSO.

We next investigated whether lower doses of AAV2-retro can yield robust retrograde transfection of ipsilateral LOC neurons while minimizing the labeling of contralateral LOC neurons. To this end, we injected 200 nl and 50 nl of virus solution representing 50% and 25% of the original 360-400nl dose. Reducing the dose of AAV2-retro still produced robust retrograde transfection of LOC neurons in the ipsilateral LSO (Figure 4 A-C), with an average of 163±74 (N=5) and 181±102 (N=5) labeled LOC neurons for the 200nl and 50nl respectively, which was similar to the 249±115 neurons (N=5) observed with the initial 400nl dose (Figure 4E, one-way ANOVA F=1.057 P=0.378). However, lowering the dose of AAV2-retro significantly reduced the number of retrogradely labeled LOC neurons in the contralateral LSO (Figure 4B-D), with an average of 105±32 (N=5) and 63±13 (N=5) labeled neurons in the contralateral LSO for the 200nl and 50nl respectively, compared to the 212±85 neurons (N=5) observed at the 400 nl dose (Figure 4F, oneway ANOVA, F= 10.28 P=0.0025 Tukey post hoc comparison 200 VS 400 P= 0.02, 50 VS 400 P=0.0023). Nevertheless, reducing the dose of AAV, did not completely eliminate the diffusion of viral particles to the CSF as ectopic transfection of cerebellar neurons (Figure 4G) and contralateral SGNs (Figure 4H) was still observed even at the lowest 50nl dose. Taken together, these results indicate that AAV2-retro maintains ipsilateral LOC transfection efficacy at lower doses and support the idea that contralateral labeling of LOC neurons is a dose-dependent consequence of viral particles diffusing through the CSF.

**Figure 4:**
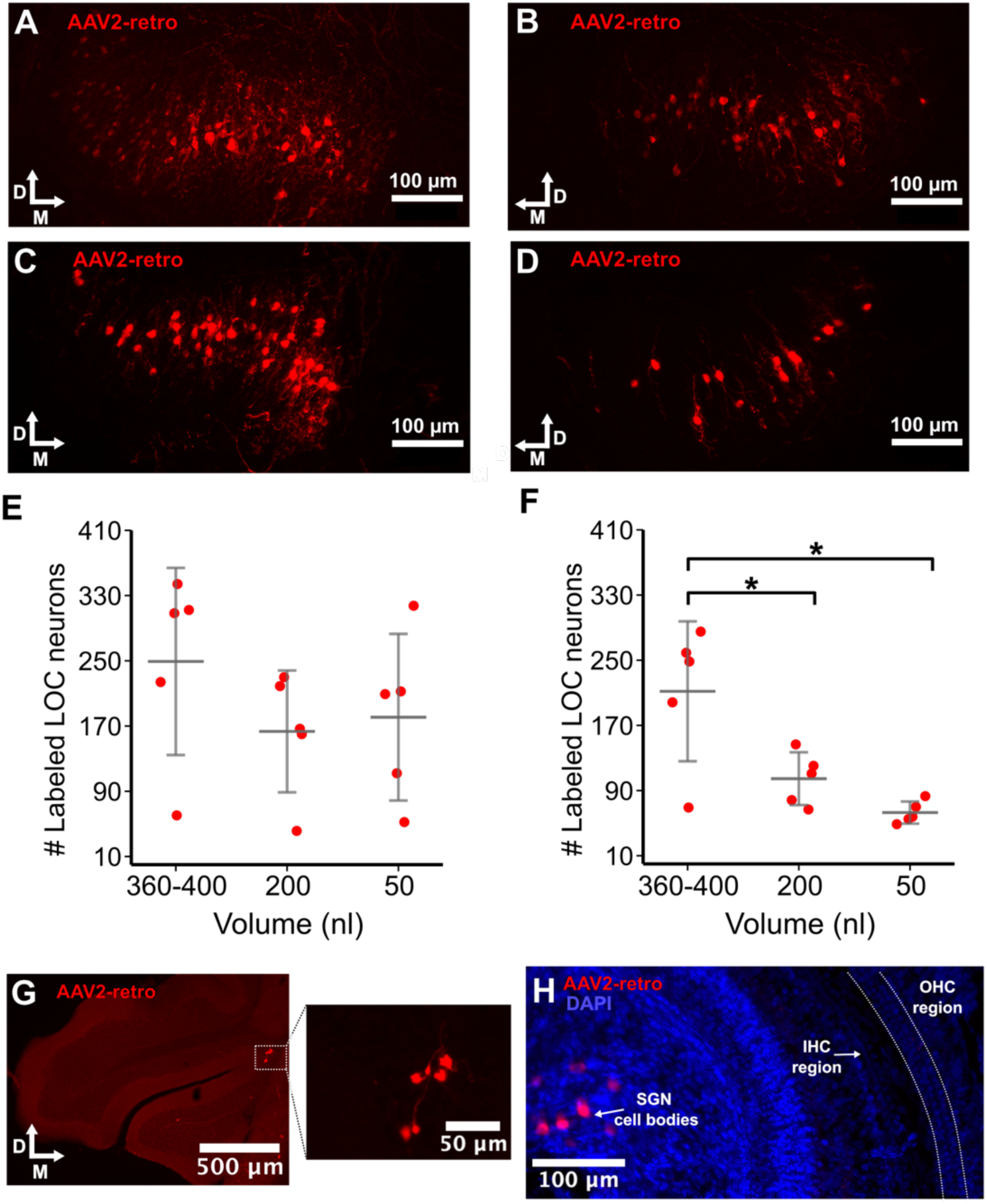
Reducing the dose of AAV2-retro still produced robust retrograde labeling of LOC neurons in the ipsilateral LSO but reduced labeling of LOC neurons in the contralateral LSO following neonatal unilateral inner ear delivery. A-B) Retrogradely labeled core LOC neurons in ipsilateral and contralateral LSO, respectively, following neonatal inner ear delivery with 200nl of AAV2-retro. M= medial, D= dorsal. C-D) Retrogradely labeled core LOC neurons in ipsilateral and contralateral LSO, respectively, following neonatal inner ear delivery with 50nl of AAV2retro. E) Quantification of retrogradely labeled core LOC neurons in ipsilateral LSO by each AAV volume injected. 360-400nl: 249±115 N=5, 200nl: 163±74 N= 5, 50nl: 181±102 N= 5. One-way ANOVA F=1.057 P=0.378. F) Quantification of retrogradely labeled core LOC neurons in contralateral LSO 360-400nl: 212±85 N=5, 200nl: 105±32 N= 5, 50nl: 63±13 N= 5; One-way ANOVA, F= 10.28 P=0.0025; Tukey post hoc comparison 200nl VS 400nl P= 0.02, 50nl VS 400nl P=0.0023, 200nl vs 50nl P=0.46. G) Transfected cerebellar neurons following neonatal inner ear delivery of 50 nl AAV2-retro. H) Contralateral cochlea following unilateral inner ear delivery of 50 nl AAV2-retro.

### Projections of core LOC neurons are exclusively ipsilateral in neonatal mice

In a variety of adult animal species 99-100% of core LOC neurons project to the ipsilateral cochlea (Campbell and Henson, 1988; Brown et al, 1993; Brown et al, 2008). However, olivocochlear projections can undergo reorganization during early development (Bruce et al. 2000; Barone et al 2019), which prompted us to investigate whether some core LOC neurons exhibit transient bilateral projections during early development, contributing to the contralateral labeling of LOC neurons we observed with AAV-retro. To avoid transfection of the contralateral cochlea by AAV2retro we used the chemical tracer Fast Blue, which has been previously used to trace olivocochlear efferents (Aschoff et al, 1988; Azeredo et al, 1999; Cantos et al, 2000; Horvath et al, 2000) and is not susceptible to off-target labeling in the contralateral cochlea. Unilateral injection of Fast Blue into the cochlea of neonatal mice resulted in 157.8± 38.7 (N=4) retrogradely labeled core LOC neurons in the ipsilateral LSO, which was comparable to the number we observed in mice that received 50nl AAV2-retro (181±102) (Figure 5A and C; t-test, t=0.46 P=0.66). However, Fast Blue labelled very few core LOC neurons in the contralateral LSO (Figure 5B) with an average of 3±1.8 (N= 4) neurons, which was substantially less than the 63±13 neurons (N=5) observed with the 50nl AAV2-retro dose (Figure 5D; t-test, t=9.79 P=0.0005). The sparse contralateral LSO labeling is not due to a failure of the tracer to label long-distance neurons because MOC neurons in the contralateral SOC were robustly labeled by Fast Blue (Figure 5B). Indeed, the number of retrogradely labeled MOC neurons in the contralateral SOC was larger than that of MOC neurons in the ipsilateral SOC at 24 ± 7.6 and 10 ± 3.9 respectively (Figure 5E; N=4, paired t-test, P=0.0035 t=6.653), which is in line with past literature reports demonstrating that the MOC pathway in mice is 63-75% contralateral (Campbell and Henson, 1988; Brown et al, 2008).

**Figure 5:**
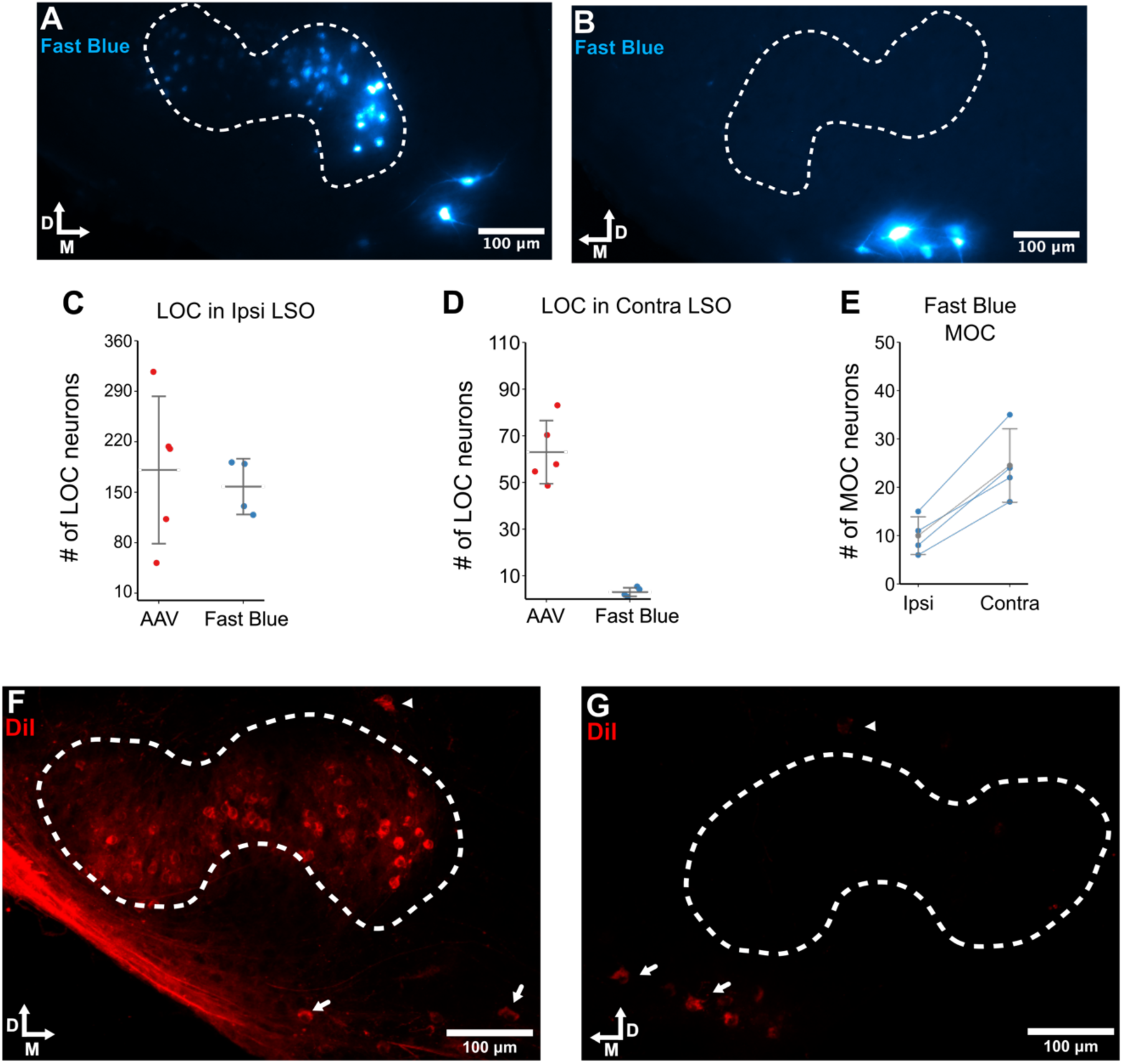
Neonatal unilateral inner ear injections of Fast Blue and DiI result in ipsilateral labeling of core LOC neurons and bilateral labeling of MOC neurons. A-B) Retrogradely labeled OC neurons in the ipsilateral and contralateral SOC, respectively, following unilateral inner ear injection of Fast Blue. M= medial, D= dorsal. C) Comparison of the number of retrogradely labeled LOC neurons in the ipsilateral LSO following 50 nl AAV2-retro and Fast Blue delivery to the inner ear. 50nl AAV2-retro: 181±102 (N=5); Fast Blue: 157.8± 38.7 (N=4); t-test, t=0.46 P=0.66. D) Comparison of the number of retrogradely labeled LOC neurons in the contralateral LSO following 50 nl AAV2-retro and Fast Blue delivery to the inner ear 50nl AAV2-retro: 63±13 (N=5); Fast Blue: 3±1.8 (N= 4); t-test, t=9.79 P=0.0005. E) Quantification of retrogradely labeled MOC neurons following Fast Blue delivery to the inner ear. Ipsilateral SOC: 24 ± 7.6; Contralateral SOC: 10 ± 3.9; (N=4) paired t-test, P=0.0035 t=6.653. F-G) Retrogradely labeled OC neurons in the ipsilateral and contralateral SOC, respectively, following unilateral DiI infusion of the inner ear. Arrowheads: Shell LOC neurons. Arrows: MOC neurons.

Fast Blue tracing requires endosomal uptake by intact axon terminals (Köbbert et al., 2000) and active axonal retrograde transport (Johnston et al 1986), which may favor the labelling of thick MOC axons over the thinner LOC axons leading to more labeling of MOC neurons in the contralateral SOC compared to LOC. We thus conducted a final set of retrograde tracing experiments using DiI, a lipophilic tracer that diffuses passively in the bilipid layer of neurons and does not depend on endosomal uptake or active axonal transport. DiI has been previously used to reliably trace the afferent and efferent innervation of the cochlea in neonatal animals (Kandler and Friauf, 1993; Bruce et al, 2000). In the LSO ipsilateral to the DiI-infused cochlea, core LOC neurons were retrogradely labeled with DiI throughout the entire length of the LSO (Figure 5F), whereas the contralateral LSO contained little to no retrogradely labeled core LOC neurons (Figure 5G). As was the case with the Fast Blue tracing experiment, the labeling of MOC neurons was robust both in the ipsilateral (Figure 5F) and contralateral (Figure 5G) SOC. Taken together, the results obtained with DiI and Fast Blue tracing indicate that the efferent projections of core LOC neurons are indeed ipsilateral in neonatal mice and that most of the contralateral LOC labeling we observed following unilateral delivery of the smallest, 50nl, dose tested can be reliably attributed to the diffusion of viral particles from the injected ear.

## Discussion

In this study, we investigated the efficacy of the AAV2-retro serotype in retrogradely transfecting core neurons of the LOC pathway. We report that injection of AAV2-retro into the posterior semicircular canal in neonatal and juvenile mice produced robust transfection of olivocochlear effect neurons in the SOC, including core LOC neurons along the entire mediolateral axis of the LSO (Figure 1B, D), which demonstrates that AAV2-retro can transfect the terminals of projecting LOC axons across all turns of the cochlea. Our results differ from those of Wang et al (2021(B)), who did not report transfected LOC neurons after AAV2-retro injections into the inner ear. The most likely reason for this apparent discrepancy is that Wang et al (2021(B)) were primarily concerned with the role of MOC and periolivary efferents, which, in the mouse, are distributed along a much wider rostro-caudal extent compared to core LOC neurons and the LSO (Brown et al 2008). It is thus possible that the brainstem sections where Wang et al (2021(B)) confirmed the presence of robust viral expression in MOC and periolivary neurons did not contain the LSO nucleus.

Unexpectedly, we also found that unilateral inner ear injection of AAV2-retro produced persistent labeling of contralateral LOC neurons (Figures 3, 4, 8). Our tracing with Fast Blue and DiI indicates that bilateral labeling of the LOC pathway was not a result of a hitherto unknown bilateral projections of LOC neurons or that it indicated a transient developmentally regulated contralateral projection, but that it stemmed from the ability of AAV2-retro to diffuse though the cochlear aqueduct to the CNS and into the contralateral cochlea. This widespread systemic diffusion of AAV2-retro particles out of the infected ear was most likely mediated by the cochlear aqueduct, which connects the CSF of the CNS to the perilymph of the cochlea (Kellerhals, 1979; Gopen et al, 1997; Talaei et al, 2019). We report that the amount of AAV2-retro delivered was directly correlated to the extent of contralateral cochlea transfection achieved via systemic viral spread throughout the CNS (Figure 4F). This observation is consistent with prior literature reports on varying levels of transfection in the contralateral cochlea following unilateral delivery of AAV. For example, György et al (2018) reported contralateral transfection in only 20% of the animals following delivery of 1-1.2 ul of AAV-PHP.B at a titer of 5-18e+10 GC/ml; whereas, studies that delivered similar volumes of AAV-PHP.B but at higher titers (1.07e+12 to 1.10e+13 GC/ml) reported observing contralateral transfection in all the animals that underwent unilateral inner ear delivery of the virus (Keppeler et al. 2018; Bali et al 2022; Mittring et al 2023). We also report that age plays an important role in determining the extent of systemic CNS spread and contralateral cochlear transfection as unilateral inner ear delivery of AAV in neonatal mice resulted in a much more robust transfection LOC efferents in the contralateral cochlea (Figure 2C-E) compared to what was observed following inner ear delivery in juvenile mice (Figure 2J-L). Our observation that AAV achieves much higher systemic spread in the neonatal CNS is in line with previous studies that delivery of AAV to the CSF in neonates results in much more widespread and robust transfection in the CNS compared to adults (von Jonquieres et al, 2013; Mathiesen et al, 2020; Garza et al, 2025).

The most important factor behind systemic viral spread throughout the CNS and onto the contralateral cochlea, however, is arguably the AAV serotype utilized. When the same titer and volume of AAV are delivered to the CSF in animals of the same age, some AAV serotypes achieve much higher rates of widespread and robust transfection of CNS neurons compared to others (Chakrabarty et al 2013; Hammond et al, 2017; Chauhan et al, 2024). Indeed, AAV serotypes that have been reported to achieve transfection of the contralateral cochlea following unilateral inner ear delivery like AAV-Anc80L65 (Landegger et al, 2017) and AAV-PHP.B (Keppeler et al. 2018; Bali et al 2022; Mittring et al 2023), have also been reported to achieve more widespread transfection of CNS neurons following delivery to the CSF compared to other serotypes (Deverman et al, 2016; Hudry et al 2018; Mathiesen et al, 2020). The AAV2-retro serotype was created from AAV2 by inserting a sequence of 10 amino acids and introducing two point mutations in the VP1 capsid gene (Tervo et al, 2016). Compared to other AAV serotypes that transduce neuronal tissue, AAV2 exhibits some of the lowest levels of systemic spread throughout the CNS following delivery to the inner ear (Han et al, 2024), CSF (Passini et al, 2003; Broekman et al 2006), and bloodstream (Zincarelli et al 2008). The modification of the AAV2 capsid gene, therefore, not only enhanced the retrograde transfection efficacy of AAV2-retro, but also increased it propensity of systemic spread throughout the CNS.

AAV2-retro exhibits an attenuated binding affinity for heparan sulfate proteoglycans (HSPG) compared to its ancestral serotype, AAV2 (Tervo et al, 2016). HSPGs can be broadly classified into transmembrane HSPGs, like syndecan and glypican, that are found on the cell surface and secreted HSPGs, like perlecan and agrin, that are present in the extracellular matrix (ECM) (Condomitti et al, 2018; Kamimura et al, 2021; Matsuzaka et al, 2024). The strong HSPG binding affinity of AAV2 (Summerford et al 1998; Opie et al 2003; Perablo et al 2006; Lochrie et al 2006; O’Donnell et al 2008) leads to its sequestration by ECM-bound HSPGs, which in turn limits its ability to spread from its injection site (Cehajic-Kapetanovic et al, 2011). Accordingly, attenuation of the the HSPG binding affinity enhances increases the spread of AAV2 from its injection site (Gorbatyuk et al, 2019). The high expression of ECM-bound HSPGs in cochlear basement membrane tissue (Torihara et al, 1994; Cosgrove et al, 1996; Tsupurn et al, 2001) coupled with the high affinity that AAV2 has for HSPG could explain why, compared to other serotypes, AAV2 produces some of the lowest levels of systematic spread throughout the CNS following unilateral inner ear delivery (Han et al, 2024). Conversely, the attenuated HSPG binding affinity of AAV2retro likely prevented its sequestration to the ECM of the injected cochlea’s tissue and contributed to its ability to transfect neurons throughout the CNS and to robustly transfect SGN cell bodies and LOC efferent terminals in the contralateral cochlea (Figures 2, 3 and 4).

In addition to the difference in their propensity for systemic spread throughout the CNS, AAV2retro and its ancestral serotype, AAV2, also seem to differ when it comes to their cellular tropism in the cochlea. Previous studies have shown that delivery of AAV2 to the inner ear results in the transfection of inner and outer hair cells (Kilpatrick et al, 2011; Iranfar et al, 2025). However, we report that AAV2-retro failed to transfect either hair cell type, as it primarily transfected efferent terminals and SGN cell bodies in the cochlea (Figure 3D). Even though AAV2 achieves cellular entry through interacting with the receptor AAVR (Pillay et al 2016; Meyer et al 2020), the HSPG binding affinity of AAV2 can influence its tropism through determining the extent of its sequestration on the surface of cells with rich expression of HSPGs (Perablo et al, 2006). Indeed, attenuating the HSPG binding affinity of AAV2 reduces its transfection efficacy of these cells (Summerford et al 1998; Opie et al 2003; Perablo et al 2006; Lochrie et al 2006). HSPG expression in cochlear hair cells plays an important role in development (Freeman et al 2014) and has been proposed to underlie the differences in viral transfection efficacy between outer and inner hair cells (Iranfar et al, 2025). It is possible, therefore, that the attenuated HSPG binding affinity of AAV2retro contributed to its reduced cellular tropism for hair cells. It is worth noting, however, that AAV serotypes which exhibit weak HSPG binding affinity, like AAV8 (Opie et al, 2003; Wu et al, 2006), were also successfully transfect hair cells (Kilpatrick et al, 2011). As such, while the attenuated HSPG binding affinity of AAV2-retro may have contributed to its inability to transfect hair cells, it is very unlikely that HSPG binding affinity was the only factor behind the change in the cellular tropism of AAV2-retro.

Taken together, our results indicate that the AAV2-retro is an effective serotype for retrograde gene delivery to core LOC neurons. We report, however, that AAV2-retro differs from its ancestral serotype, AAV2, along other properties beyond its enhanced capacity for retrograde transfection. Most consequential of these differences was that the inner ear delivery of AAV2-retro resulted in a level of systemic spread throughout the CNS and contralateral cochlea that far exceeded what has been reported for AAV2 (Han et al, 2024). Insufficient consideration of the ability of AAV2retro to spread far from its injection site could, therefore, result in erroneous interpretation of results. For example, AAV2-retro delivery to the hippocampus was reported by Bian et al (2019) to label anterior cingulate area (ACA) efferent projections to the hippocampus; however, follow up studies that utilized AAV2 (Glat et al, 2022) and AAV2/9 (Shi et al, 2022) as anterograde tracers did not observe these ACA to hippocampus projections. Finally, Andrianova et al (2023) demonstrated through the use of Fast Blue and meticulous analysis of AAV2-retro transfection pattern that the previous report of ACA efferent projection to the hippocampus (Bian et al, 2019) was almost certainly caused by off-target AAV2-retro transfection. In conclusion, it is imperative to carefully consider the propensity of AAV2-retro to spread far from its injection site when planning experiments in order to utilize a titer and volume that minimizes the risk of off-target labeling. Additionally, our study contributes to a growing body of literature highlighting the need of careful checking any unexpected projections discovered using AAV2-retro, and other even more virulent retrograde AAV serotypes (Lin et al, 2020; Han et al 2023), through the use of a second, preferably chemical, tracing approach.

## Author Contributions

K.K. and M. K. designed the research. M.K. and M.B. performed research and analyzed data. M.K. wrote the paper. K.K. edited the paper.

## Acknowledgements

We thank Brian Brockway and Ethan Herring for help with immunohistochemistry. We thank members of the Kandler lab for their valuable discussion.

